# Biological control protects tropical forests

**DOI:** 10.1101/391748

**Authors:** K.A.G. Wyckhuys, A.C. Hughes, C. Buamas, A.C. Johnson, L. Vasseur, L. Reymondin, J.P. Deguine, D. Sheil

**Affiliations:** Zhejiang University, Hangzhou 310000, People’s Republic of China; University of Queensland, Brisbane, Australia; Joint International Research Laboratory of Ecological Pest Control, Ministry of Education, Fuzhou 350002, People’s Republic of China; China Academy of Agricultural Sciences CAAS, Beijing, People’s Republic of China; CGIAR Program on Roots, Tubers and Banana (CRP-RTB), International Center for Tropical Agriculture CIAT, Hanoi, Viet Nam; Xishuangbanna Tropical Botanical Gardens, China Academy of Sciences, Xishuangbanna (Yunnan), China; Department of Agriculture (DoA), Ministry of Agriculture and Cooperatives, Bangkok, Thailand; Graham Centre for Agricultural Innovation, Charles Sturt University, Orange, NSW, Australia; Brock University, St. Catharines (Ontario), Canada; International Center for Tropical Agriculture CIAT, Hanoi, Viet Nam; CIRAD, La Reunion, France; Norwegian University of Life Sciences, P.O. Box 5003, 1432 Ås, Norway

## Abstract

Biological control of invasive species can restore crop yields, and thus ease land pressure and contribute to forest conservation. In this study, we show how biological control against the mealybug *Phenacoccus manihoti* (Hemiptera) slowed deforestation across Southeast Asia. In Thailand, the newly-arrived mealybug caused an 18% decline in cassava yields over 2009-2010, a shortfall in national production and an escalation in the price of cassava products. This spurred an expansion of cassava cropping in neighboring countries from 713,000 ha in 2009 to >1 million ha by 2011: satellite imagery reveal 388%, 330%, 185% and 608% increases in peak deforestation rates in Cambodia, Lao PDR, Myanmar and Viet Nam focused in cassava crop expansion areas. Following release of the host-specific natural enemy *Anagyrus lopezi* (Hymenoptera) in 2010, mealybug outbreaks were reduced, cropping area contracted and associated deforestation slowed by 31-94% in individual countries. When used with due caution and according to current guidelines, biological control offers broad benefits for people and the environment.

---

The UN Sustainable Development Goals aim to end malnutrition and poverty while also preventing biodiversity loss^1^. These goals place competing demands on land that are not readily^2-4^. reconciled. For example, agricultural expansion serves many fundamental needs but often results in the clearing of forests with negative consequences for biodiversity, fresh water and atmospheric composition^5,6^. Given the need to reconcile such competing demands on land, we must identify and promote all appropriate options including those, such as arthropod biological control, that are often disregarded. Here, for the first time, we show how biological control can reduce pressure on land and thus spare forests.

Invasive species, including many agricultural pests, constrain the production of food and other commodities ^7^, and often impose additional costs such as the disruption of ecosystem services (e.g., nutrient cycling), damage to infrastructure or increased disease in humans^8^. Since the late 1800s, more than 200 invasive insect pests and over 50 weeds across the globe have been completely or partially suppressed through biological control, often with favorable benefit:cost ratios (ranging from 5:1 to >1,000:1)^9, 10^. Modern biological control, centered on a careful selection and subsequent introduction of a specialized natural enemy (from the pest species’ region of origin), thus offers an effective solution for invasive species problems ^11^. This approach is particularly useful in smallholder farming systems, as biological control is self-propagating and requires little involvement from local stakeholders^12^. Nonetheless there are risks as exemplified by few (poorly-selected) control agents that have subsequently become major problems themselves, such as the cane toad *Buffo marinus* or the weevil *Rhinocyllus conicus*^13, 14^. A consequence is that, despite significant improvements in risk assessment and management over the past three decades, concerns often obscure the potential benefits and result in biological control being avoided when it could be valuable^13^. While the failures of the last century appear well known, the more modern success stories require wider recognition. Our goal here is to present one such story.

In 2010, biological control was implemented against the invasive mealybug, *Phenacoccus manihoti* (Hemiptera: Pseudococcidae) that had first been detected in late 2008 in Thailand’s cassava crop. Grown on nearly 4 million ha across tropical Asia and extensively traded, cassava, *Manihot esculenta* Crantz, is a globally-important source of starch, a food staple for vulnerable rural populations in several Asian countries, and a base for the production of food products, animal feed, ethanol and household items^15^. In Southeast Asia, cassava is cultivated as much in small-scale diversified systems by smallholders as in large plantations. Upon arrival in Asia, *P. manihoti* spread to its ecological limits (yet confined by cassava cropping area)^16^, leading to an average 4.1 ton/ha reduction in crop yield in Thailand (from 22.7 to 18.6 ton/ha), a 27% drop in the nation’s aggregate cassava production and an ensuing 162% increase in starch price^15^. One response was the 2009 introduction of the host-specific parasitoid wasp *Anagyrus lopezi* (Hymenoptera: Encyrtidae; originally native to South America) from Benin (West Africa), where it had suppressed *P. manihoti* throughout Africa following its introduction in 1981 ^17^. These wasps were released across Thailand from mid-2010 onward, and were subsequently introduced into Lao PDR, Cambodia and Viet Nam (2011-2013). They established successfully and suppressed mealybug populations across the region^18^. This restored yields by 5.3-10.0 tonnes/ha (as assessed through manipulative assays), and helped stabilize the trade in cassava root, starch and substitute commodities (i.e., maize, wheat, potato)^15^.

In this study, we characterized how the cassava mealybug invasion and ensuing biological control are associated with agricultural expansion and forest loss in mainland Southeast Asia. These forests include the most species-rich and biologically-valuable habitats in the region^19,20^. We first conducted surveys to quantify the extent of parasitoid-mediated *P. manihoti* population suppression (*section i*). Second, we examined regional cassava cultivation and trade from 2009 to 2013 (*section ii*). Third, we contrasted forest loss and cassava expansion over the period (*section iii*).

## Results

### i. Regional pest & parasitoid survey

Our surveys, conducted across mainland Southeast Asia between 2014 to 2017 (i.e., 6-9 years and 5-8 years following the initial *P. manihoti* detection and *A. lopezi* introduction, respectively), showed that *P. manihoti* was present in 37.0% of the fields (*n* = 549) and comprised 20.8% abundance within a speciose mealybug complex (Fig. 1). Among sites, *P. manihoti* reached field-level incidence of 7.6 ± 15.9% (mean ± SD; i.e., proportion mealybug-affected tips) and abundance of 5.2 ± 19.8 insects per infested tip. *Anagyrus lopezi* wasps were recorded in 96.9% of mealybug-affected fields (*n* = 97), at highly-variable parasitism rates. For example, in mid- to large-scale plantations parasitism rates ranged from 10.7 ± 10.6% (*n* = 20; Dong Nai, Vietnam) to 67.1 ± 20.8% (*n* = 22) in late dry season in Tay Ninh (Vietnam). In low-input, smallholder-managed systems (see methods), parasitism varied between 17.1 ± 14.8% (*n* = 18; Ba Ria Vung Tau – BRVT, Vietnam) to 46.7 ± 27.8% in central Cambodia (*n* = 10). Where *A. lopezi* was present, mealybug abundance was negatively associated with *A. lopezi* parasitism (ANOVA, F 1,84= 12.615, p= 0.001; Fig. 1;^18^).

**Figure 1.**
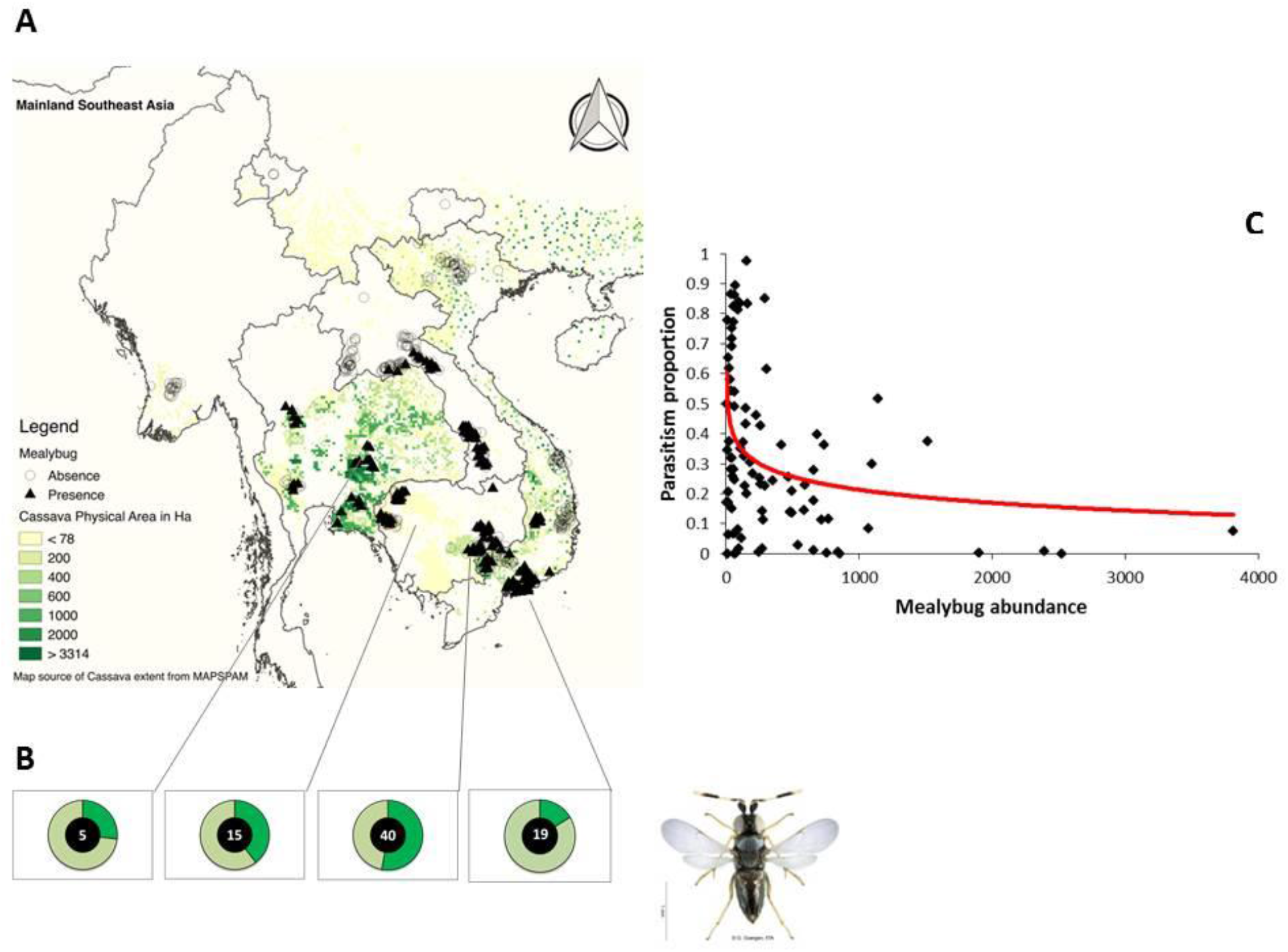
Map of Southeast Asia depicting ***P. manihoti*** geographical distribution, complemented with field-level ***A. lopezi*** parasitism and mealybug abundance records. In *panel A*, green shading reflects the approx. 4 million ha of cassava cultivated regionally in 2005 (MapSpam, 2017). *Panel B* presents doughnut charts, indicative of the percent *A. lopezi* parasitism (as depicted by the dark green section) at four selected sites. The number inside each doughnut reflects the number of fields sampled per locale. *Panel C* presents the relationship between average *P. manihoti* abundance and *A. lopezi* parasitism level per field, for a total of 90 fields in which simultaneous recordings were done of mealybug infestation pressure and parasitism rate (figure adapted from Wyckhuys et al., 2018).

### ii. Country-specific cassava production and trade

In Thailand, cassava cropping area reached 1.32 million ha in 2009, and subsequently fell to 1.18 million (2010) and 1.13 million ha (2011). This followed the country-wide *P. manihoti* outbreak in 2009, yield losses as experienced over an 8-10 month time lapse (with cassava a semi-perennial crop, routinely harvested at 8-10 months of age), and reduced total cassava production. Over the ensuing 2009-10 cropping season, province-level yields dropped by 12.59 ± 9.78% (area-weighted mean: −18.2%) and country-wide aggregate yields declined from 22.67 t/ha to 18.57 t/ha (Fig. 2). Regional production followed similar trends: total production in Viet Nam, Myanmar, Lao PDR and Cambodia dropped from 66.93 million tonnes in 2009 (at yields of 19.42 t/ha) to 62.04 million tonnes in 2010 (at 18.56 t/ha). Yet, over 2009-2011, 129.7%, 387.0%, 52.8%, and 16.0% increases were recorded in the volume of harvested cassava root in Cambodia, Lao PDR, Myanmar and Viet Nam respectively. Over this period, cassava cropping area in all countries increased substantially, for example expanding from 160,326 ha to 369,518 ha in Cambodia (Fig. 3; Supplementary Figure 1).

**Figure 2.**
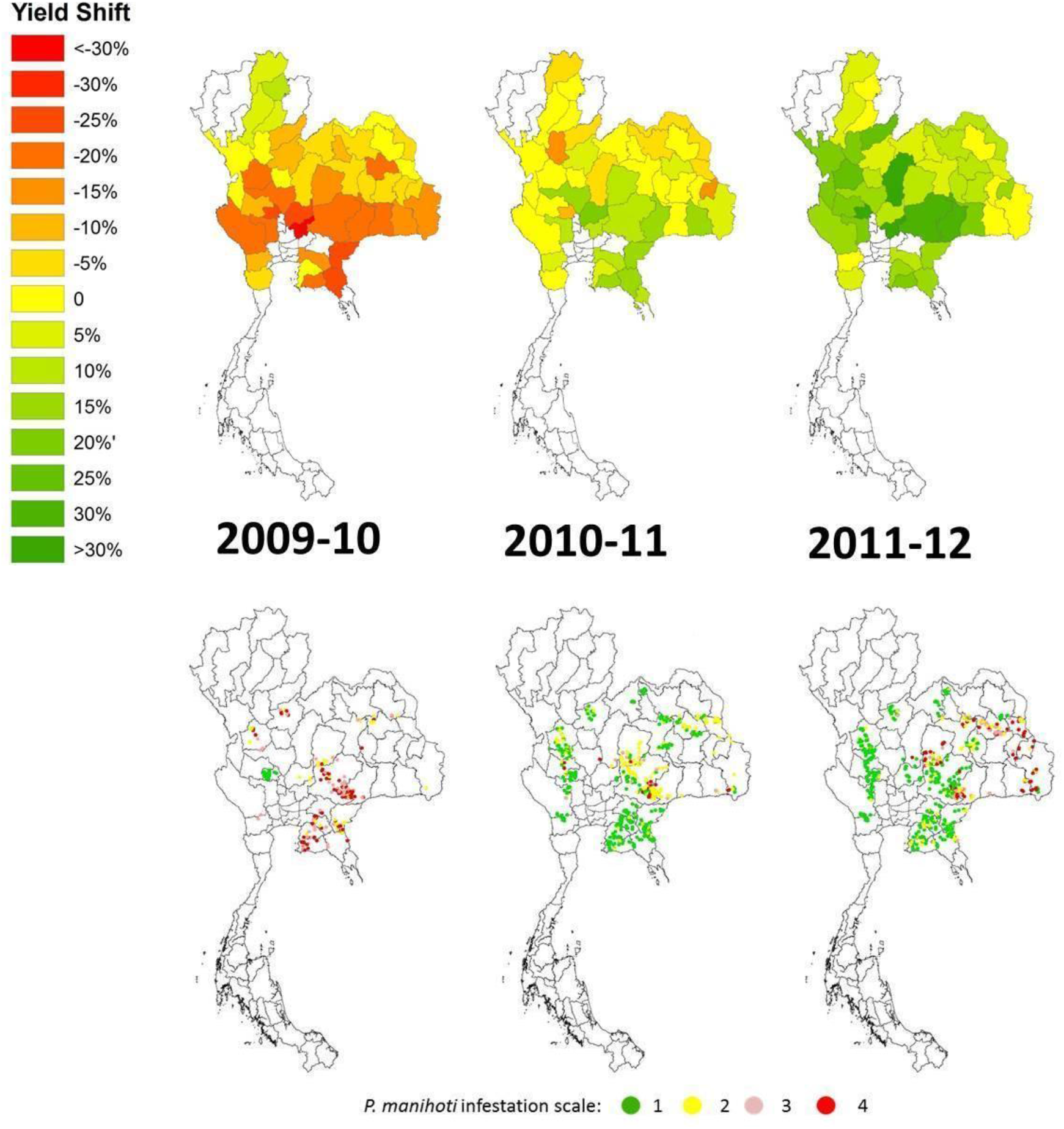
Yield recovery following biological control in Thailand’s cassava crop over 2009-12. Patterns are reflective of the country-wide cassava mealybug invasion (late 2008 onward) and ensuing biological control campaign. The *upper panel* reflects annual change in cassava crop yield (for a given year, in percent as compared to the previous year) for a select set of provinces (adapted from Wyckhuys et al., 2018). In the *lower panel,* historical records of *P. manihoti* spatial distribution and field-level infestation pressure are shown over successive growing seasons (data facilitated through Thai Royal Government).

**Figure 3.**
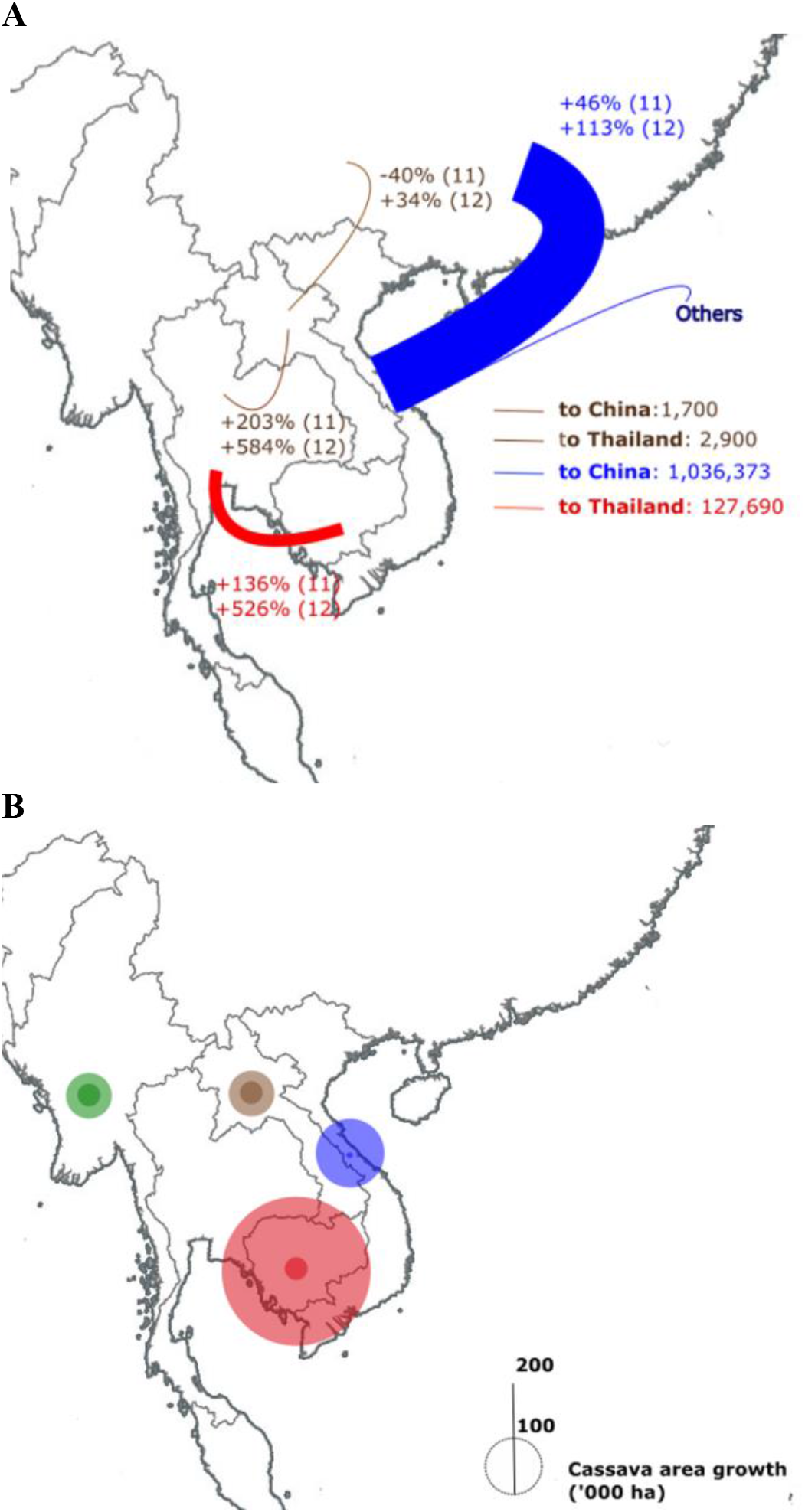
Annual shifts in inter-country cassava trade mirror country-level expansion of cassava cropping area. In *panel A,* export volume is depicted of cassava roots, chips and pellets from Cambodia and Lao PDR (to Thailand and China) and Vietnam (to China). Thickness of the arrow reflects relative volume of traded cassava, and yearly increases in export volume are specified for 2010-2011, and 2011-2012 (A). *PanelB* depicts the annual rate of increase in harvested cassava area (ha) for individual Southeast Asian countries (except Thailand), from 2009 until 2011.

From 2009 to 2012 regional trade in cassava-based commodities shifted, as Thailand’s import of cassava products (i.e., roots, chips and pellets) increased by 153% and starch by 1,575%, and Viet Nam exported more of these products to China. In 2009, Thai imports represented 1,126 tonnes of cassava products from Lao PDR, and 322,889 tonnes from Cambodia. By 2012, respective volumes had attained 19,844 tonnes and 799,456 tonnes. In 2009, Viet Nam’s exports of cassava products to China reached 2.09 million tonnes, and increased to 2.21 million tonnes by 2012. Over this period there was a regional increase in cassava cropping area from 713,000 ha (2009) to >1.02 million ha by 2011 (Fig. 3; Supplementary Figure 1). In all countries except Lao PDR, cropping area was greatest in 2011 and reached 369,500 ha in Cambodia, 56,500 ha in Myanmar, and 558,200 ha in Viet Nam (Supplementary Fig. 2). By 2013, cassava area contracted and Thailand’s import trade of cassava products and cassava starch lowered by a respective 42.3% and 83.5%.

### iii. Country-specific forest loss vs. cassava area

Regional deforestation surged in 2010 with an annual net loss of 653,500 ha from 278,900 during the preceding year (Terra-i; Fig. 4), partially mirroring the 302,000 ha increase in cassava area harvested during 2011 (for an 8-10 month-long crop; see above). Over 2010, Terra-i estimated total forest loss to be 166,700 ha in Cambodia, 157,600 ha in Viet Nam, 74,700 ha in Lao PDR, and 254,400 ha in Myanmar; a respective 169%, 207%, 80% and 104% higher than in 2009. Between 2006 and 2012, the first months of 2010 represented the peak of deforestation and reached 20,181 ha/week in Cambodia, 17,015 ha/week in Viet Nam and 51,284 ha/week in Myanmar. For Lao PDR, a peak of 10,128 ha/week was attained in early 2011 (Supplementary Fig. 2). Peak deforestation rates during the 2010 dry season were a respective 388%, 608%, 185% higher than those in 2009, and for Lao PDR represented a 330% increase. By 2011, peak deforestation rates in Cambodia, Viet Nam and Myanmar lowered with a respective 42.0%, 31.8% and 94.9% compared to 2010, while peak deforestation rates in Lao PDR lowered by 50.5% in 2012.

**Figure 4.**
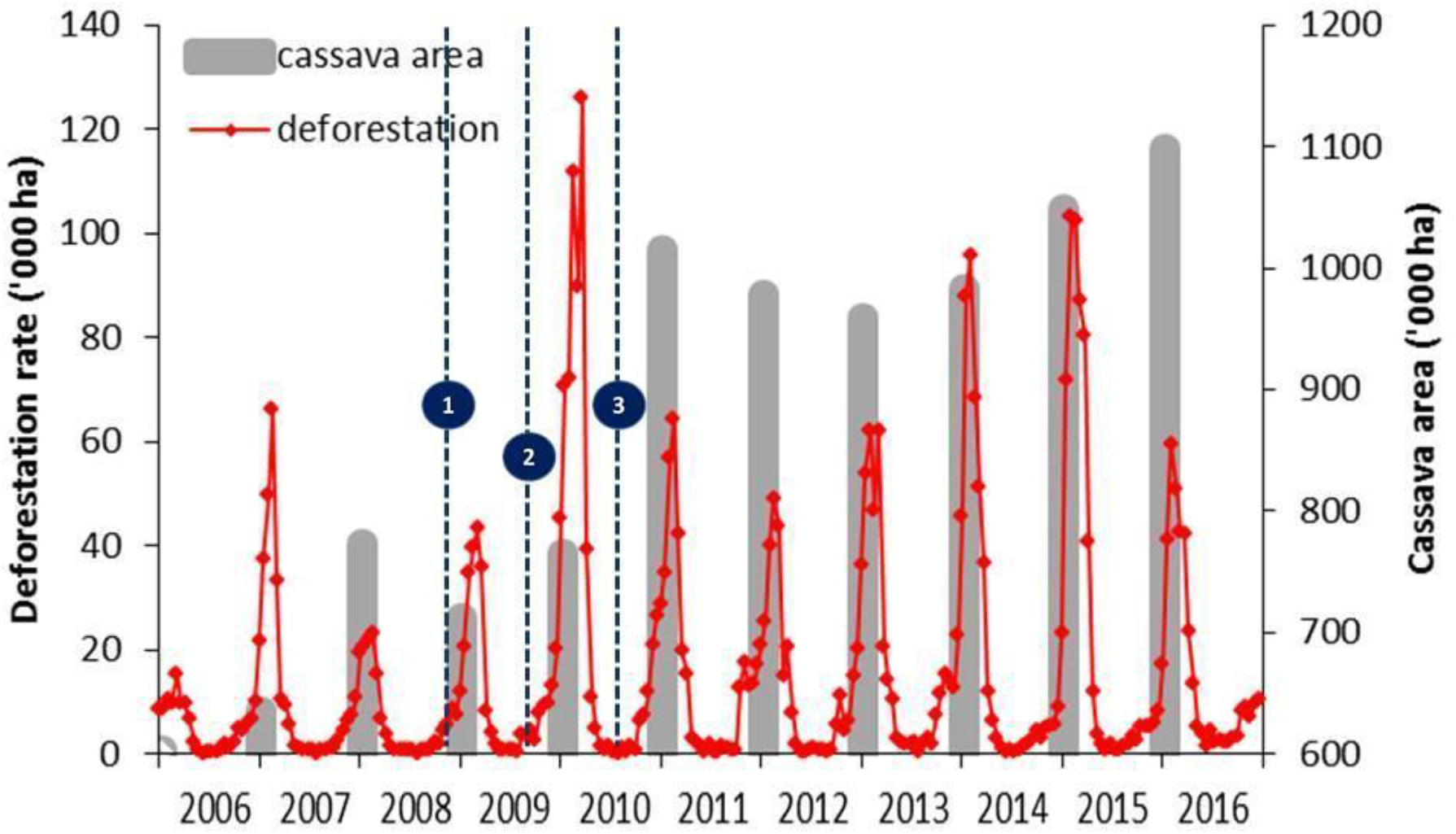
Near real-time deforestation patterns relate to the annual increase in (harvested) cassava area over a 2006-2016 time period. Patterns cover a) the late 2008 invasion and subsequent regional spread of *P. manihoti* (event # 1), b) the initial introduction of *A. lopezi* from Benin, West Africa (event #2), and c) nation-wide parasitoid release in cassava fields across Thailand (event # 3). Deforestation and cassava area growth statistics are compiled from individual country records of Lao PDR, Vietnam, Cambodia and Myanmar.

Examining patterns for provinces, a significant association was recorded between (province-level, summed) regional deforestation and cassava area growth over 2005-2010 (ANOVA; F_1_,_60_= 14.278, p< 0.001), over 2010-2013 (F_1,54_= 18.240, p< 0.001) and over the entire 2005-2013 time period (F_1_,_65_= 18.011, p< 0.001) (Fig. 5, 6). For Viet Nam specifically, province-level forest loss was positively related to extent of (harvested) cassava area growth during 2011-2012 (F_1_,_24_= 7.113, p= 0.013) and 2012-2013 (F_1_,_20_= 4.603, p= 0.044), but not during 2009-2010 (F_1_,_27_= 0.295, p= 0.591) or 2010-2011 (F_1_,_40_= 2.863, p= 0.098). Similar patterns and associations were recorded for Cambodia for 2005-2010 and 2010-2012 (Supplementary Fig. 3). Since 2014, deforestation in Cambodia and Viet Nam has continued (Supplementary Fig. 2), likely reflecting growth of China’s demand for cassava products.

**Figure 5.**
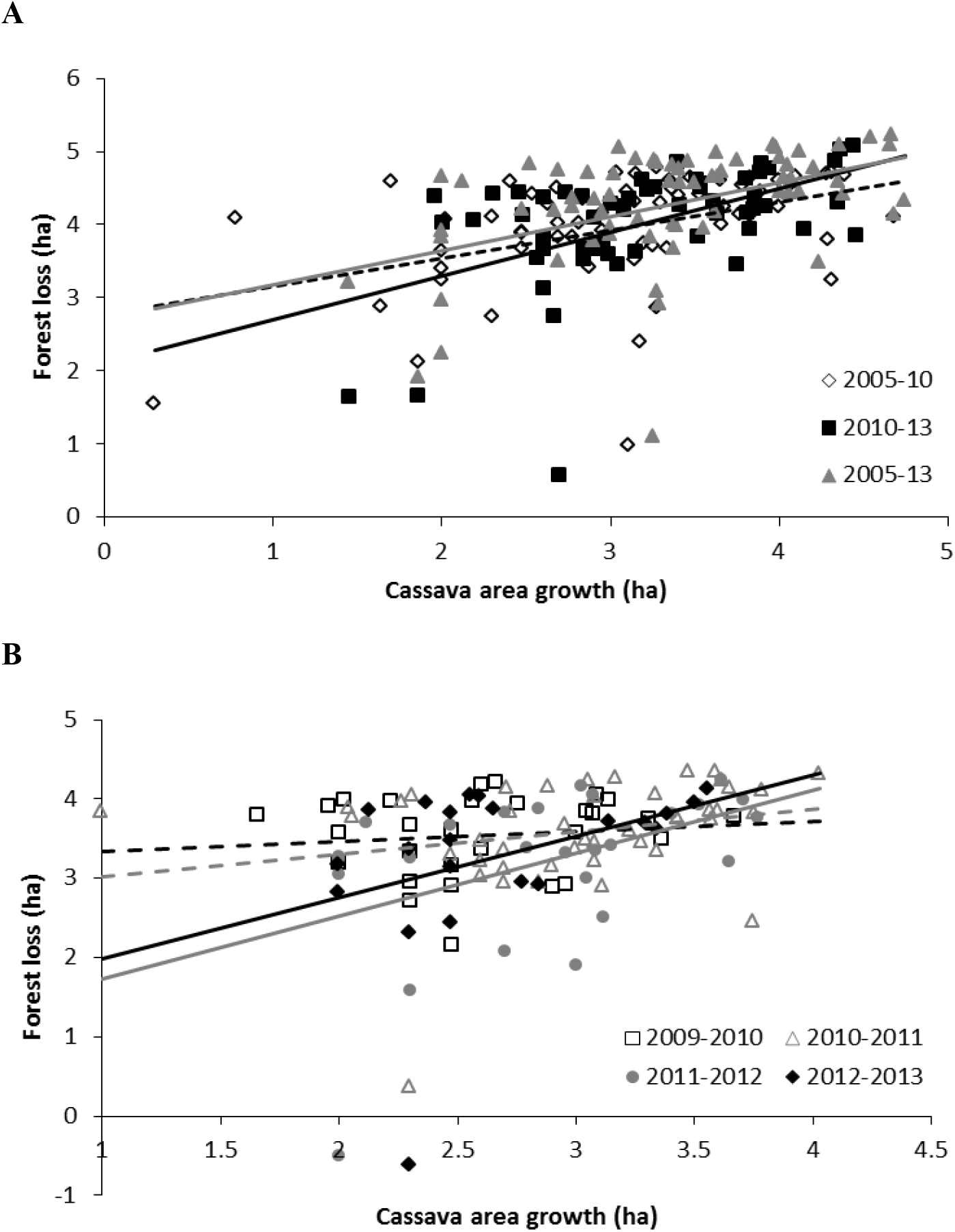
Regional and country-specific patterns in deforestation relate to growth of cassava cropping area over 2005-2013. *Panel A* represents regional patterns, showing province-level cassava area increase (ha) in Viet Nam, Cambodia and Lao PDR as related to degree of forest loss (ha) over a 2005-10, 2010-2013 and entire 2005-2013 time frame. *Panel B* contrasts annual forest loss against increase in (harvested) cassava area, for 40 different Vietnamese provinces. Both variables are log-transformed, and only certain regression lines in *panelB* reflect statistically significant patterns (ANOVA, p< 0.05; see text for further statistics). Data are exclusively shown for provinces and time-periods in which cassava area expansion was recorded. Dashed lines represent patterns for 2005-10 (panel A) and 2009-10, 2010-11 (panel B).

**Figure 6.**
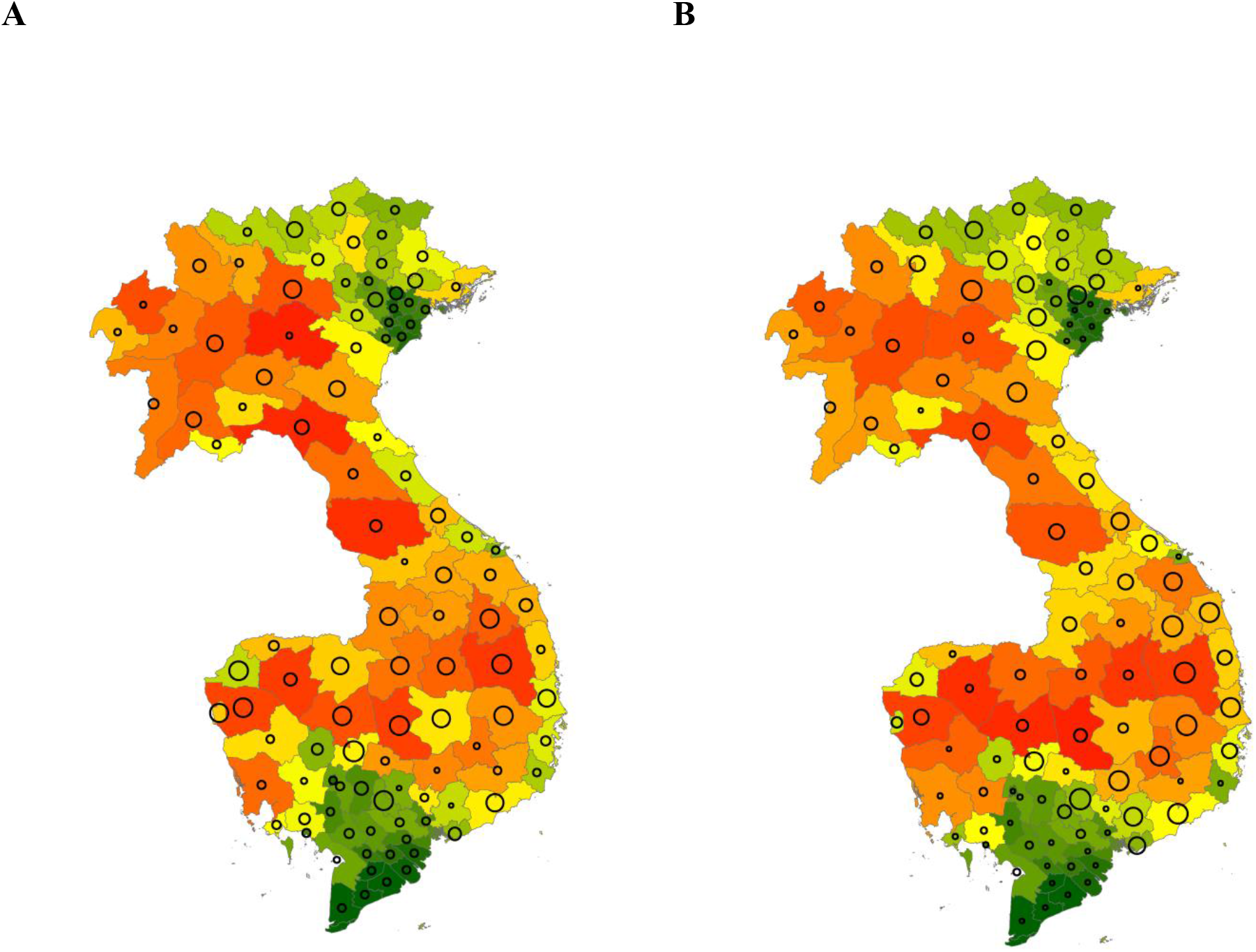
Forest loss relates to cassava area expansion across the Greater Mekong subregion, over two distinct time periods (i.e., 2005-2010, A; 2010-2013, B). Province-level deforestation and cassava area growth over particular time periods are contrasted for Lao PDR, Cambodia and Vietnam, with bubble size depicting cassava area growth (ha) and coloring reflecting level of forest loss (with increasing levels of forest loss indicated by colors ranging from green to red).

## Discussion

We have shown how the 2008 *Phenacoccus manihoti* invasion in Thailand contributed to a >300,000 ha increase in cassava cropping area in neighboring countries, to make up for a shortfall in supply and an 138-162% surge in cassava prices. More specifically, as the mealybug invasion is associated with a 243-281% increase in imports of cassava products, cassava trade thus likely became a key regional driver of deforestation. The mealybug invasion also prompted broad and recurrent use of insecticides in Thailand (Supplementary files; Supplementary Fig. 4), with potential impacts on biodiversity, human health^21^ and ecosystem functioning^22,23^, including interference with *P. manihoti* biological control (Wyckhuys et al., unpublished). Given the importance of cassava for local smallholder families, the changes in cassava productivity (e.g., total crop loss in 2009-2010 in parts of Thailand and western Cambodia) also had marked socioeconomic impacts including declines in farmer income. The introduction of *A. lopezi*, allowed Thailand’s cassava production to recover, helped stabilize cassava trade, averted the need for insecticides in neighboring countries and reduced cassava-driven forest loss in the region.

Demand for cassava is an important driver of land-use pressure and forest loss in the Greater Mekong sub-region, yet it is not the only one, and the *A. lopezi* introduction alone thus will not avert future deforestation. Other drivers of forest loss are the establishment of oil palm, pulp and paper plantations, rubber and food crops^24^; crops that are cultivated across tropical Asia through significant engagement from agro-enterprises^24,25^, with their actions regularly affected by ‘place-less’ incentives (e.g., varietal improvements)^26,27^, (foreign-based) consumer demand^15^, or chronic soil fertility loss e.g., for cassava in fragile upland settings^28^. Yet, during 2010-2012, our analyses revealed the marked role of cassava area growth in deforestation at a multi-country level. To stabilize the forest margin, a multi-functional ‘landscape approach’ and a systematic analysis of the various drivers of land-use change will be necessary^29^. Also, in order to enhance the capacity of cropping systems to absorb (or recover from) perturbances such as the *P. manihoti* attack, indices can be adopted that reflect broader ‘ecosystem resilience’. Through use of those indices, agro-industry can simultaneously contribute to agricultural sustainability and biodiversity conservation^30,31^. Furthermore, such ‘resilience’ indices could be employed by local government and agro-industry actors alike to further encourage good practice. By stabilizing cassava yields and alleviating pressure on land and dependence on insecticides, biological control supports agricultural intensification and spares land for conservation^4, 32^. Nonetheless, while such land-sparing activities are valuable, these are insufficient to achieve conservation in the long-term without suitable policies, planning, governance arrangements, funding and implementation^29,33^.

Several factors contributed to the success of the mealybug biological control program. These include: early detection (e.g.,^34^); proper identification of the pest^35^; availability of and unrestricted access to an effective host-specific parasitoid^36^ and decisive action with private-sector involvement. These factors allowed an effective program to be swiftly planned, assessed and implemented^37,38^, without the benefits of biological control being obscured by the risks. Though few cases have justifiably blemished the reputation of arthropod biological control, current practices and safeguards minimize such risks^13,39^. Our study equally helps put those risks into perspective, as the rapid *A. lopezi* introduction proved essential to alleviate the disruptive impacts of *P. manihoti* attack^34,40^. The benefits gained through the A. lopezi release thus need to be viewed in light of the multi-faceted ecosystem impacts of invasive species^41,42^, and the environmentally-disruptive actions that are regularly employed for their mitigation^43,44^. Our study illustrates how an invasive pest can lead to substantial loss of forest^45^ and accelerating species loss and extinctions^24,46^, while biological control offered a powerful, environmentally-benign tool to permanently resolve those impacts^11^. Now, by concurrently highlighting the harmful and beneficial impacts of *P. manihoti* and *A. lopezi* respectively, our work can enable a concerted search for ‘win-win’ solutions that address invasive species mitigation, biodiversity conservation and profitable farming.

Biological control requires a reassessment by all those responsible for achieving a better world ^2–4, 47, 48^. While invasive species undermine many of the UN Sustainable Development Goals^1, 8, 49^ the benefits of biological control are routinely disregarded^13,48^. Though an objective appraisal of risks remains essential, an equivalent recognition of the benefits is also warranted. When used with established safeguards^13^, biological control can resolve or reduce the problems caused by invasive species^11^ and helps ensure crop protection benefits not only farmers^38^, but also the environment.

## Methods

### i. Pest & parasitoid survey

Insects were surveyed in 549 cassava fields in Myanmar, Thailand, Lao PDR, Cambodia, Viet Nam and southern China, from early 2014 until mid-2015, using standard protocols (see^18^). Briefly, 8-10 month old fields in the main cassava-growing provinces of each country were selected with assistance from local plant health authorities, with sites located at least 1 km apart and within easy reach by vehicle. Surveys were conducted January-May 2014 (dry season), October-November 2014 (late rainy season) and January-March 2015 (dry season). Locations were recorded using a handheld GPS (Garmin Ltd, Olathe, KS). Five linear transects were established per field (or site), departing from positions along an X sampling pattern covering the entire cassava field. Ten consecutive plants were sampled along each transect, thus yielding a total of 50 plants per site. Each plant was assessed for the presence and abundance (i.e., number of individuals per infested tip) of *P. manihoti*. In-field identification of *P. manihoti* was based on morphological characters, and samples were equally transferred to the laboratory for further taxonomic confirmation. For each site, average *P. manihoti* abundance (per infested tip) and field-level incidence (i.e., proportion of *P. manihoti-infested* tips) was calculated.

To contrast local *P. manihoti* infestation pressure with *A. lopezi* parasitism rates, we sampled during 2014 and 2015 at a random sub-set of mealybug-invaded sites in different provinces in Thailand (*n* = 5), Cambodia (n = 10, 15 per province), and southern Vietnam (*n* = 18, 20, 22). In doing so, samples were obtained from both smallholder-managed, diversified systems (i.e., 1-2 ha in size) and from mid- to large-scale plantations (i.e., at least 5-10 ha in size). Sampling for *A. lopezi* parasitism consisted of collecting 20 mealybug-infested cassava plant tips at each site which were transferred to a field laboratory for subsequent parasitoid emergence. Upon arrival in the laboratory, each cassava plant tip was examined, predators were removed and *P. manihoti* counted. Next, tips were placed singly into transparent polyvinyl chloride (PVC) containers, covered with fine cotton mesh. Over the course of three weeks, containers were inspected daily for emergence of *A. lopezi* parasitic wasps. Parasitism levels of *A. lopezi* (per tip and per site) were calculated. Next, for sites where *A. lopezi* was found, we analyzed field-level *P. manihoti* abundance with *A. lopezi* parasitism rate with linear regression (see also^18^). Variables were log-transformed to meet assumptions of normality and homoscedasticity, and all statistical analyses were conducted using SPSS (PASW Statistics 18).

### ii. Country-specific cassava production and trade trends

To assess how mealybug invasion and ensuing parasitoid-mediated cassava yield recovery affected cassava production and trade, we examined country-level production and inter-country trade for cassava-derived commodities. More specifically, we contrasted cassava yield and production trends with inter-country trade flows over periods spanning the 2008 *P. manihoti* invasion, the 2009 *A. lopezi* introduction into Thailand and the subsequent (natural, and human-aided) continent-wide distribution of *A. lopezi* (mid-2010 onward). Our assessments detailed shifts in cassava production (harvested area, ha) and yearly trade flows (quantity) of cassava-derived commodities into Thailand from neighboring countries within the *P. manihoti* invaded range.

Crop production statistics for Thailand were obtained through the Office of Agricultural Economics (OAE), Ministry of Agriculture & Cooperatives (Bangkok, Thailand). Furthermore, country-specific patterns of cassava production (harvested area, ha) and yield (t/ha) were obtained for Viet Nam, Myanmar, Lao PDR and Cambodia via the FAO STAT database (http://www.fao.org/faostat/). To assess structural changes in the inter-country trade of cassava-derived commodities, we extracted data from the United Nations Comtrade database (https://comtrade.un.org/). Over a 2006-2016 time period, we recorded the following evolutions in terms of quantity (tonnes): global annual imports of cassava-derived commodities to Thailand (reporting) and China, from ‘All’ trade partner countries. More specifically, we queried the database for bilateral trade records of three cassava-derived commodities and associated Harmonized System (HS) codes: “Cassava whether or not sliced - as pellets, fresh or dried” (71410), “Tapioca & substitutes prepared from starch” (1903), and “Cassava starch” (110814). Given the occasional inconsistencies in country-reported trade volumes or values in either FAO STAT or Comtrade databases, cross-checks were made with databases from the Thai Tapioca Starch Association (TTSA) and rectifications were made accordingly.

### iii. Country-specific trends in forest loss vs. cassava area growth

To infer the likely impact of cassava area growth on forest loss in different Southeast Asian countries, we obtained data from both a near-real time vegetation monitoring system, Terra-i (https://www.terra-i.org) and deforestation data from Global Forest Watch^50^ (https://www.globalforestwatch.org/). Terra-i relies upon satellite-derived rainfall and vegetation data obtained through TRMM sensor data (Tropical Rainfall Monitoring Mission) and MODIS MOD13Q1 respectively, to detect deviations from natural vegetation phenology patterns that cannot be explained by climatic events. More specifically, Terra-i adopts computational neural networks to detect how vegetation vigor behaves at a given site over a period of time in relation to observed rainfall, and thus identifies certain anomalies while accounting for the effects of drought, flooding and cloud cover or other image ‘noise’. Changes in vegetation greenness at the landscape level are recorded on the Terra-i platform on a bi-weekly basis. Terra-i outputs have been validated through comparison with the Global Forest Change data and the PRODES system in Brazil. This showed that these datasets are similar as the average KAPPA coefficient was of 0. 96 ± 0.004. Furthermore, the average recall value for detection of events with an area of 90% to 100% of a MODIS pixel is of 0.9 ± 0.05 which shows that Terra-i detects the large-size events. However, an average recall of 0.28± 0.13 has been observed when the event size is about 10% of a MODIS pixel, showing a limitation of Terra-I to detect smaller size tree cover clearance. Country-level deforestation statistics over a 10-year time period were extracted from this platform for Lao PDR, Myanmar, Viet Nam and Cambodia, and data were compiled on a province level for each year from 2005 to 2013.

Next, yearly province-level records of cassava (harvested) area were compiled for each of the different countries by accessing FAO STAT, the Cambodia 2013 agriculture census and primary datasets as facilitated through national authorities and the International Food Policy Research Institute (IFPRI), Washington DC, USA. For Lao PDR, province-level records were only available on cultivated area of all root crops combined. Here, we assumed that major inter-annual changes in harvested area of root crops in Lao PDR can be ascribed to cassava as other locally-important root crops, such as yam and sweetpotato are mostly grown for subsistence purposes and are less subject to major inter-annual area shifts. No continuous yearly datasets on local cassava cultivation area were available for Cambodia, and no province-level cassava cultivation records could be accessed for Myanmar. Because of these variations in available data some analyses were carried out over different periods (see below).

To quantify the extent to which forest loss was related to cassava area expansion, two types of analyses were conducted. First, we used linear regression to relate province-level increases in harvested cassava cropping area with forest loss during that same period for all countries (i.e., Cambodia, Lao PDR, Viet Nam), over three different time frames: 2005-2010, 2010-2013 and 2005-2013. Second, as complete annual records on (province-level) cassava cultivation were available for Viet Nam, linear regression analysis allowed annual province-level trends in forest loss to be related to cassava expansion for individual years (2009-2013) relating. To meet assumptions of normality and heteroscedasticity, data were subject to log-normal (for cassava area records) or rank-based inversed normal transformation (for deforestation rates and records). All statistical analyses were conducted using SPSS (PASW Statistics 18)

### iv. Data Availability

Upon acceptance of the manuscript, all data will be made available in an appropriate public structured data depository, and the accession number(s) provided in the manuscript.

## Supporting information

**Supplementary Figure 2.**
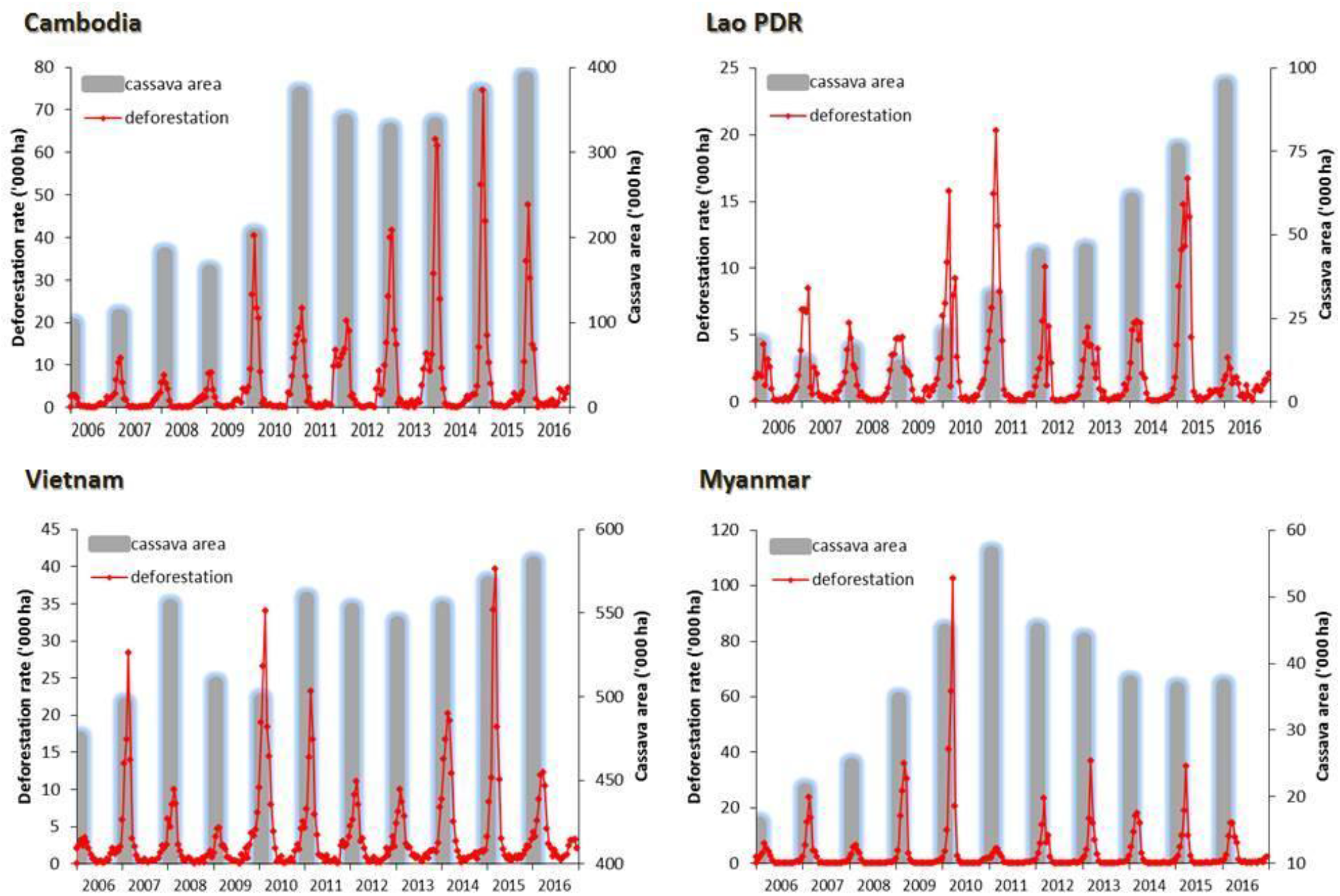
Country-specific deforestation patterns as related to the annual increase in (harvested) cassava area over a 2006-2016 time period, covering the late 2008 invasion and subsequent continent-wide spread of *P. manihoti*, and the release of *A. lopezi* across Thailand in mid-2010.

**Supplementary Figure 3.**
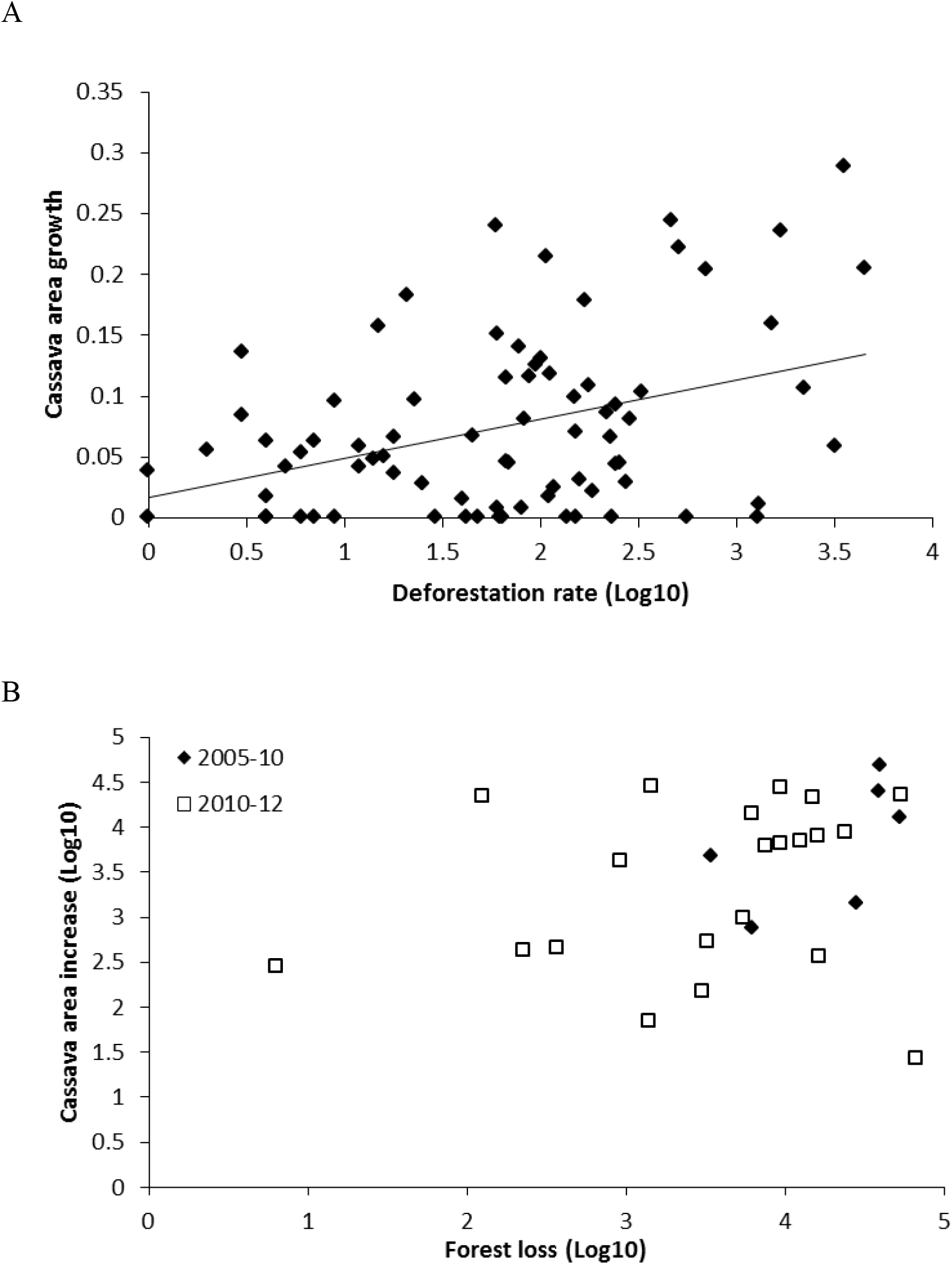
Country-level cassava area-growth patterns, as related to local forest loss. Panel A depicts the proportional annual increase in (planted) cassava area as related to local degree of forest loss during the preceding year, for 40 different Vietnamese provinces over a 2008-2012 time period. *Panel B* represents province-level cassava area increase (ha) as related to degree of forest loss (ha), for 24 Cambodian provinces over a 2005-10 and 2010-12 time frame. Data are exclusively shown for provinces and time-periods in which cassava area expanded. Annual deforestation rates are log-transformed, and the regression line in *panel A* reflects a statistically significant pattern (ANOVA, p< 0.05).

## Farmer adoption of insecticide use

### 1. Materials & Methods

From 2014 until 2016, extensive farmer surveys were conducted in Thailand, Cambodia and Vietnam. In all three countries, household-level surveys were carried out using semi-structured questionnaires with open-ended questions, to optimally gauge farmer’s knowledge and pest management behavior. One interview was done per household, following a person-to-person interview format. A semi-structured questionnaire was employed, with open-ended questions to better elicit farmer knowledge. The questionnaire was pre-tested and revised prior to use at the national level, in a given country. In Thailand, surveys were entirely carried out by local officers from the Thai Department of Agricultural Extension (DoAE), while trained enumerators supported by local authorities (General Directorate of Agriculture of Cambodia, and Plant Protection Department, Vietnam) took part in survey activities in the other two countries. At all sites, surveys were conducted by interviewers that were fluent in the local languages. Though survey instruments were designed for multiple purposes (e.g., Delaquis et al., unpublished), we only cover pest management activities in this study. For assessment of local pest management behavior, farmers were asked to freely enumerate knowledge and adoption of management practices for control of *P. manihoti.* Particular attention was paid to farmers’ reported usage of (preventative) dips with neonicotinoid insecticides.

In Thailand, farmer surveys were conducted in a total of 33 cassava-growing provinces over the course of 2014 (i.e., 6 years after the initial detection of *P. manihoti).* In each province, a variable number of farmers was interviewed by DoAE personnel, ranging from n= 20 (Roy-et, Payao) to n= 348 (Karnchanaburi), attaining a grand total of 2,505 cassava farmers in the national territory. Sample size was determined by local authorities, and is only partially reflective of the number of cassava growers in a given province. In Cambodia and Vietnam, nation-wide surveys were carried out in a more systematic fashion in at least 15 districts per country during late 2016 (i.e., 8 years following the initial detection of *P. manihoti* in Thailand). In these two countries, interviews were focused in districts with the largest area of cassava production. Within each district, a total of 15 cassava growers were randomly selected and interviewed, attaining the following respective sample sizes for Cambodia and Vietnam: n= 240 (16 districts) and n= 206 (15 districts). District-level adoption rates were pooled per province for either country, and mapped at a regional scale.

### 2. Results

In Thailand, 71.3% farmers (*n* = 2,505) used prophylactic dips with systemic insecticides for P. manihoti management (Supplementary Fig. 3). Regional adoption rates of insecticide dips ranged from 45.8% in eastern parts to 90.3% in northern areas of the country. Province-level rates of insecticide use were highest in Payao and Tak (100%; n= 20, 38 respectively), Lampang (98.0%; n= 49), Utaradit (96.1%; n= 26), Yasothorn (92.9%; n= 84) and Loei (91.9%; n= 123).

**Supplementary Figure 4.**
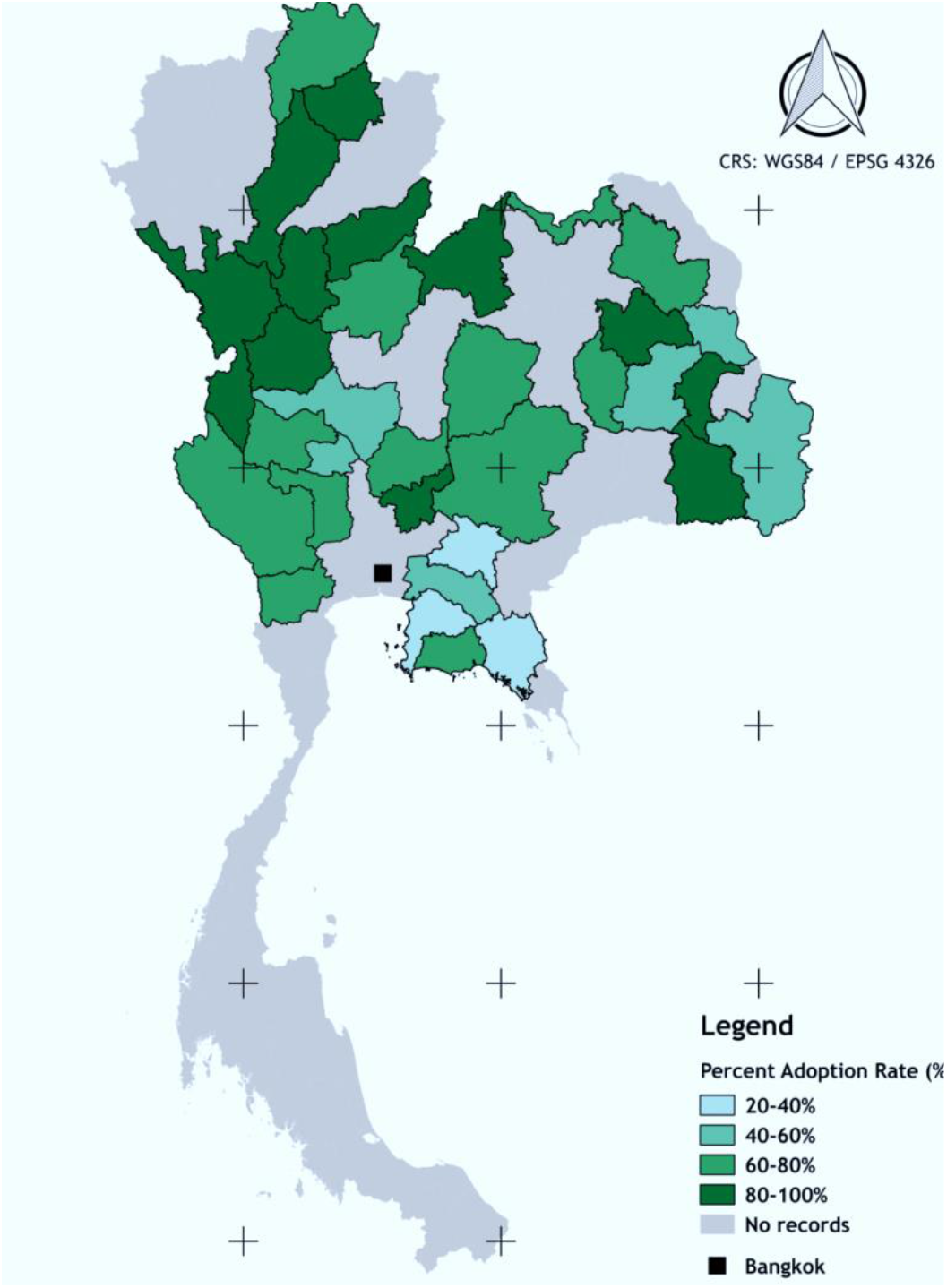
Level of adoption of prophylactic insecticide dips amongst cassava farmers in 33 different provinces across Thailand, as recorded during mid-2014. Adoption levels are expressed as % of surveyed farmers in each province, within a nationwide survey of 2,500 growers (*n* = 20-348 per province). For provinces in grey, no data were obtained.

